# Index-based map-to-sequence alignment in large eukaryotic genomes

**DOI:** 10.1101/017194

**Authors:** Davide Verzotto, Audrey S.M. Teo, Axel M. Hillmer, Niranjan Nagarajan

## Abstract

Resolution of complex repeat structures and rearrangements in the assembly and analysis of large eukaryotic genomes is often aided by a combination of high-throughput sequencing and mapping technologies (e.g. optical restriction mapping). In particular, mapping technologies can generate sparse maps of large DNA fragments (150 kbp–2 Mbp) and thus provide a unique source of information for disambiguating complex rearrangements in cancer genomes. Despite their utility, combining high-throughput sequencing and mapping technologies has been challenging due to the lack of efficient and freely available software for robustly aligning maps to sequences. Here we introduce two new map-to-sequence alignment algorithms that efficiently and accurately align high-throughput mapping datasets to large, eukaryotic genomes while accounting for high error rates. In order to do so, these methods (OPTIMA for glocal and OPTIMA-Overlap for overlap alignment) exploit the ability to create efficient data structures that index continuous-valued mapping data while accounting for errors. We also introduce an approach for evaluating the significance of alignments that avoids expensive permutation-based tests while being agnostic to technology-dependent error rates. Our benchmarking results suggest that OPTIMA and OPTIMA-Overlap outperform state-of-the-art approaches in sensitivity (1.6–2× improvement) while simultaneously being more efficient (170–200%) and precise in their alignments (99% precision). These advantages are independent of the quality of the data, suggesting that our indexing approach and statistical evaluation are robust and provide improved sensitivity while guaranteeing high precision.

## 1. Introduction

In recent years, the availability of commercial platforms for high-throughput genome mapping (e.g. from OpGen, BioNano Genomics, and Nabsys) have increased the interest in using these technologies, in combination with high-throughput sequencing data, for applications such as structural variation analysis and genome assembly. In particular, several recent genome assembly projects have highlighted their utility for obtaining high-quality assemblies of large eukaryotic genomes (e.g. for goat [7] and budgerigar [8] genomes) or studying complex genomic regions [11] and cancer genomes [15]. Mapping technologies typically provide sparse information (an ordered enumeration of fragment sizes between consecutive genomic patterns, e.g. restriction sites) for very large fragments of DNA (150 kbp–2 Mbp) and are thus orthogonal in utility to sequencing approaches that provide a base-pair level information for smaller fragments. Combining these two pieces of information therefore requires effective algorithms to align maps to sequences.

Alignment of maps (typically called Rmaps, for restriction maps) to sequences has been widely studied as an algorithmic problem, with a range of practical applications: from genome scaffolding [14] to assembly improvement [12] and validation [5]. The general approach to do so has been to translate sequence data to get *in silico* maps and comparing these to experimentally obtained maps using dynamic programming algorithms. For large genomes and mapping datasets, naive all-versus-all dynamic programming can be computationally expensive. On the other hand, high error rates in mapping data (e.g. optical mapping can miss 1 in 4 restriction sites) has led to the use of model-based scoring functions for sensitively evaluating alignments [17,1,16]. These often require prior knowledge and modeling of mapping error rates (e.g. fragment sizing errors, false cuts and missing cuts) and can be expensive to compute [2,3,17]. Alternative approaches, with simpler (non-model-based) scoring functions [14] are handicapped by the need to do expensive permutation-based statistical testing to evaluate the significance of alignments. While these approaches work well for microbial genomes, they typically do not scale well for larger genomes, where they might also have limited sensitivity. In contrast, commercially available solutions for map-to-sequence alignment (e.g. Genome-Builder from OpGen) scale better and have been used for the assembly of large eukaryotic genomes [7], but tend to discard a large fraction of the mapping data (>90%) due to reduced sensitivity and correspondingly lead to increased mapping costs for a project.

Map-to-sequence alignment algorithms are thus faced with the twin challenges of improving sensitivity and precision on one end, and reducing computational costs for alignment and statistical evaluation on the other end. An elegant solution to this problem from the field of sequence-to-sequence alignment is the use of a seed-and-extend approach [9]. However, since maps represent ordered lists of continuous values, this extension is not straightforward, particularly when multiple sources of mapping error and their high error rates are taken into account [13]. In addition, since error rates can vary across technologies, and even across different runs on the same machine, it is not clear if a general and sensitive map-to-sequence aligner is feasible. An efficient statistical testing framework that helps control for false discovery without *a priori* information about error rates is indeed critical for making such an aligner easy to use and applicable across technology platforms.

In this work, we describe how a sorted search index and the use of a composite seeding strategy can help efficiently and sensitively detect seed map-to-sequence alignments. Our second contribution is in the design of a robust and fast statistical evaluation approach that includes the contribution of multiple sources of mapping errors in the alignment score and evaluates the significance of the best alignment using all identified, feasible solutions. We incorporated these ideas as well as additional refinements to solve two common alignment problems: glocal alignment, with OPTIMA, where an entire map is aligned to a subsequence of a second (typically *in silico*) map, and overlap alignment, with OPTIMA-Overlap, where the end of one map is aligned to the beginning of another. When benchmarked against state-of-the-art aligners, OPTIMA and OPTIMA-Overlap typically provide a strong boost in sensitivity (1.6–2×) without sacrificing precision of alignments (about 99%). Moreover, our pilot implementations exhibited runtime improvements over commercially available tools (2× over OpGen’s Gentig) and orders-of-magnitude over published, freely available algorithms and software [17,14]. Finally, these methods exhibited robustness to variations in error distributions, while being agnostic to them, suggesting that they can deal with different experimental outcomes of the same technology (e.g. different map cards or lane types) as well as being applicable across mapping technologies (with minor modifications for preprocessing of data). As glocal and overlap alignments form the basis of a range of applications that involve the combination of sequence and mapping data (e.g. assembly scaffolding, refinement and validation, structural variation analysis, and resolving complex genomic regions), OPTIMA and OPTIMA-Overlap can serve as building blocks for these applications, allowing for more time and cost-effective analyses.

## 2 Definitions

High-throughput genome mapping technologies typically work by linearizing large molecules of DNA, e.g. in nanochannels [11], and using enzymes such as restriction enzymes to recognize and label (e.g. by cutting DNA) specific patterns throughout the genome, e.g. a 6-mer motif. These patterns are then read out (typically, optically) to obtain an ordered set of fragment sizes for each DNA molecule. If corresponding genome sequences or assemblies are available, these can be converted into *in silico* maps through pattern recognition [2]. Let *o*_1_, *o*_2_, …,*o*_*m*_ be the *m* ordered fragment sizes of an experimentally derived map *o*, and *r*_1_, *r*_2_, …, *r*_*n*_ be the *n* fragment sizes of an *in silico* map *r*. For simplicity we suppose here that *m* ≤ *n*. In an idealized case, we may define the problem of glocally aligning *o* to *r* as a one-to-one correspondence between all the fragments of *o* with a subset of the fragments of *r*, i.e. *r*_*l*_, *r*_*l+1*_, …, *r*_*l*+*m*−1_ (we could also reverse the roles of *o* and *r* here). In practice, many sources of errors affect experimentally derived maps including missing cuts, false/extra cuts, missing fragments, and fragment sizing errors [17]. *In silico* maps could also be affected by sequencing or assembly errors [14], but these are less likely to impact alignments as they are typically infrequent. To accommodate for errors, we extend the definition of correspondence between map fragments to allow for matches between sets of fragments (see Figure 1a), as used similarly in [14]:

**Figure 1:**
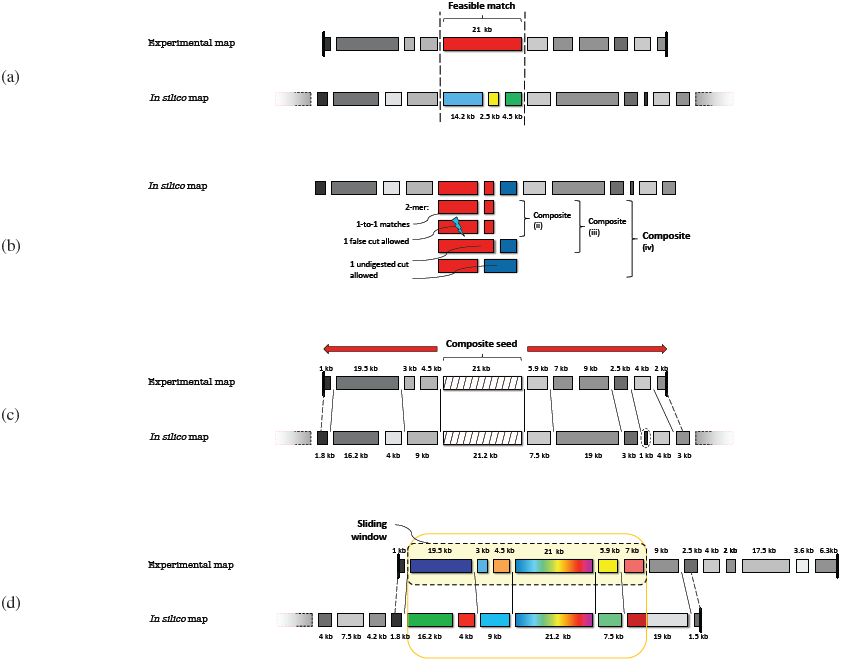
Examples of (a) *feasible match* within dashed bars (Definition 1); (b) *composite seeds* with *c* = 2 (Definition 2), where *Composite* represents the final composition of seeds with errors used here; the case with one false cut allowed is not directly indexed from the *in silico* maps, but is effectively used later in the seeding process; (c) seed extension in glocal alignment with dynamic programming (straight lines delimit feasible matches found, dashed lines mark truncated ends matches, and dashed circles show possible missing fragments); and (d) sliding window approach in overlap alignment: for a particular window of fixed size (dashed black border) we first compute a glocal alignment (solid yellow border) from one of its seeds (multicolored box), statistically evaluate it, and subsequently extend it until the end of one of the maps is reached on both sides of the seed.

### Definition 1 (Feasible match)

*A subset of fragments o_k_, o_k+l_, …, o_s_ aligned as a whole entity to a subset of in* silico fragments *r_l_, r_l+1_, …, r_t_ is said to be a feasible match if*

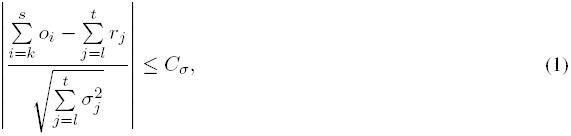

*where the σ_j_ are the standard deviations allowed for each (reference) in silico fragment size in order to match the experimental fragments, and C_σ_ = 3 is an appropriate bound if sizing errors are approximately normally distributed.*

A valid **glocal alignment** is then an ordered set of feasible matches *M*_1_, *M*_2_, …, *M*_*w*_ between experimental and *in silico* fragments, such that all the experimental fragments *o*_1_, *o*_2_, …, *o*_*m*_ are aligned to a subset of the *in silico* fragments *r*_*t*_, *r*_*t*+1_, …, *r*_*v*_, and both sets are orderly partitioned by all the matches *M*_*1…w*_ without overlaps, with *w≤m* and *w≤v−t* + 1. Missing fragments, which usually arise from short fragments below the experimental detection limit (e.g. 2 kbp), can be handled in this framework by allowing the option of ignoring short fragments for the purpose of the *C*_*σ*_ bound (Equation (1)). We next define a valid **overlap alignment** *M*_1_, *M*_2_, …, *M*_*w*_ as one that allows experimental maps and *in silico* maps to only partially align with each other, with both *M*_1_ and *M*_*w*_ corresponding to an end of one of the maps (see Figure 1d). In general, as maps can have truncated ends, we relax the *C*_*σ*_ test to be only an upper bound on sizes for experimental maps, or a lower bound for *in silico* maps, when map ends are considered.

## 3 Glocal map-to-sequence alignment

OPTIMA is the first alignment tool based on the seed-and-extend paradigm that is able to deal with erroneous mapping data. The basic paradigm is similar to that used for the alignment of discrete-valued sequences (allowing for mismatches and indels), and is as follows. We start by indexing the *in silico* maps in order to be able to smartly use this information later, and find seeds for each experimental map *o* corresponding to some indexed regions of those sequences. We then extend these seeds by using dynamic programming in order to try to align the whole experimental map to the corresponding *in silico* map region. For each map *o*, *feasible* solutions —as defined by the index structure, size of the genome, and maximum error rate— are then evaluated by a scoring scheme to select the optimal solution. Finally, the statistical significance and uniqueness of optimal solutions is determined by comparison and modeling of all the feasible solutions found.

### Continuous-valued composite seeds

The definition of appropriate seeds is critical in a seed-and-extend approach to maintain a good balance between sensitivity and speed. A direct extension of discrete-valued seeds to continuous-valued is to consider values that are close to each other (as defined by the *C*_*σ*_ bound) as being matches. However, as mapping data typically have high error rates [2,1,16] and represent short sequences (e.g. on average optical maps contain 10–22 fragments, representing roughly a 250 kbp region of the genome), a seed of *c* consecutive fragments is likely to have low sensitivity unless using a naive *c* = 1 approach (see Figure 2 for a comparison) —the latter could nevertheless easily require to visit *O*(*n*) seeds (where *n* is the total length of the *in silico* maps) as in global search approaches [1,16]. Therefore, analogous to the work on spaced seeds for discrete-valued sequences [6], we propose and validate the following composite seed extension for continuous-valued seeds:

**Figure 2.**
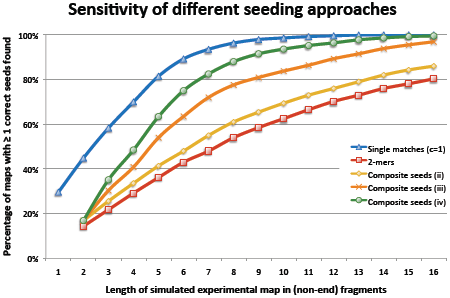
Comparison of sensitivity between different seeding approaches over a set of 1,600 simulated experimental maps (scenario (B) presented in Section 5). For each corresponding length in fragments, we report the percentage of maps with at least one correct seed detected (i.e. correct location in the *in silico* maps). For example, 88% of maps with 8 (non-end) fragments had at least one correct seed matched using our composite seeds (*Composite seeds (iv)*, as defined in Definition 2 and shown in Figure 1b).

#### Definition 2 *(Composite seeds)*

*Let r_j_*_1_, *r*_*j*2_ *and r*_*j*3_ *be consecutive restriction fragments from a reference* in silico *map. A composite seed, for c = 2, is given by including all of the following:*

i. *the c-mer r_j_*_1_, *r*_*j*2_, *corresponding to no false cuts in the* in silico *map,*
ii. *the c-mer r_j_*_1_ + *r*_*j*2_, *r*_*j*3_, *corresponding to a missing cut in the experimental map (or false cut in the in silico map), and*
iii. *the c-mer r_j_*_1_, *r*_*j*2_ + *r*_*j*3_, *corresponding to a different missing cut in the experimental map (or false cut in the* in silico *map),*

*as depicted in* Figure 1b.

The reference index would then contain all *c*-tuples corresponding to a composite seed as defined in Definition 2 for each location in the reference map. In addition, to account for false cuts in the experimental map, for each set of consecutive fragments *o*_*i*_1__, *o*_*i*_2__, and *o*_*i*_3__ in the experimental maps, we search for *c*-tuples of the type *o*_*i*_1__, *o*_*i*_2__ and *o*_*i*_1__ + *o*_*i*_2__, *o*_*i*_3__ in the index (i.e. *Composite seeds (iv)* in Figure 1b). As shown in Section 5, this approach significantly reduces the space of candidate alignments without impacting the sensitivity of the search (see also Figure 2). To index the seeds, we adopt a straightforward approach where all *c*-tuples are collected and sorted into the same index in lexicographic order (say, for the *c*_*i*_ elements, with 1 *≤ i ≤*_*c*_, in the *c*-tuple) by *c*_1_. Lookups can be performed by binary search over fragment-size intervals that satisfy the *C*_*σ*_ bound for *c*_1_, and a subsequent linear scan of the other elements *c*_*i*_, for *i ≥* 2, of the selected tuples, while verifying the *C_σ_* bound in each case. Note that, as seeds are typically expected to be of higher quality, we can apply a more stringent threshold on seed fragment size matches (e.g. we used 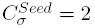).

Overall, the computational cost of finding seeds using this approach is *O*(*m* (log *n* + *c #seeds_c_*_=1_)) per experimental map, where *n* is the total length of the *in silico* maps in fragments, *m* ≪ *n* is the length of the experimental map, and *#seeds_c_*_=1_ is the number of seeds found in the first level of the index lookup, before narrowing down the list to the actual number of seeds that will be extended (i.e. #*seeds*). The cost and space of creating the reference index is thus *O*(*cn*), if the number of errors considered in the composite seeds is limited and bounded (as in Definition 2), using radix sort to sort the index. This approach drastically reduces the number of alignments computed in comparison to more general, global alignment searches [14], as will be shown later in Section 5.

### Dynamic programming-based extension of seeds

In order to extend a seed to get a glocal alignment we adopt a scoring scheme similar to that used in SOMA [14]. This allows us to evaluate alignments without relying on a Likelihood-based framework that requires prior information on error distributions as input [17]. In addition, we can use dynamic programming to efficiently find glocal alignments that optimize this score and contain the seed (Figure 1c). Specifically, we proceed along the dynamic programming matrix by aligning the end of the s-th experimental fragment with the end of t-th *in silico* fragment using backtracking to find feasible matches, i.e. those that satisfy Equation (1) and minimize the total number of cut errors (i.e. missing cuts + false cuts + missing fragments found), with ties being broken by minimizing a χ^2^ function for fragment sizing errors:

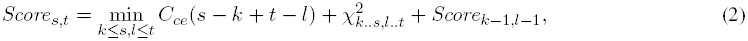

where the first index of each subscript represents experimental fragments and the second index the *in silico* fragments, *s − k* is the number of false cuts, *t − l* is the number of missing cuts, *C*_*ce*_ is a constant larger than the maximum possible total for 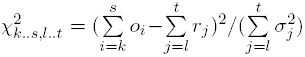 and *Score*_0;0_ = 0, else *Score*_i;0_ = ∞ and *Score*_0,j_= ∞. Similarly to [2] we band the dynamic programming and its backtracking to avoid unnecessary computation using the same *δ*-parameters. In addition, we stop the dynamic programming-based extension if no feasible solutions can be found for the current seed after having analyzed at least a number of fragments (e.g. 5) of the experimental map.

### Statistical significance and uniqueness of alignments

In order to evaluate the statistical significance of a candidate alignment we exploit the fact that we have explored the space of feasible alignments in our search, and use these alignments to approximate a random sample from a (conservative) null model. Specifically, for each candidate alignment found, we compute its distance from the null model in a feature space (to be defined later) using a Z-score transformation, and then use this score to evaluate if it is optimal, statistically significant, and unique.

We start by identifying a set *F* of orthogonal features, with respect to random alignments, that are expected to follow the Normal distribution (e.g. under the law of large numbers), and compute a Z-score for each feature *f* ∈ *F*, for each candidate solution *π* ∈ Π identified through the seeding method. Each Z-score takes into account the mean and standard deviation of *f* among all candidate solutions Π found. Accounting for all considered features *f*_*i*_, with 1 ≤ *i* ≤ *k* and *k* ≥ 2, the resulting score is given by:

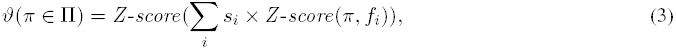

where *s*_*i*_ = 1 if lower values of feature *f*_*i*_ are preferable, and −1 otherwise, and the corresponding p-value is *p_π_* = Pnorm(*ϑ*(*π*)).

In our case, we chose a set of features based on the number of matches (indicating a higher level of conservation), the total number of cut errors, and the Wilson-Hilferty Transformation [19] of the χ^2^ score for sizing errors, WHT(*χ^2^,* #*matches*); this set can be shown to be composed of orthogonal features for false positives alignments [1,16]. The specific Z-score *ϑ*(*π*) computed for each candidate solution *π* is thus given by:

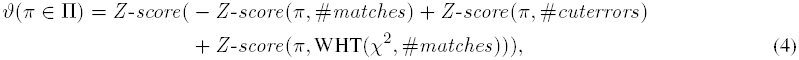

which can be subsequently converted into p-value *p*_*π*_ based on the Standard Normal distribution. The candidate solution *π*\* with the lowest p-value *p*\* is reported as the optimal solution. The statistical significance of each optimal solution can then be assessed through a FDR q-value analysis based on all candidate solutions found for comparable experimental maps, e.g. with the same number of fragments (we set *q* = 0.01 as a default threshold for reporting alignments). To assess uniqueness of a solution we implemented a test based on Cohen’s *d* effect size [10] using *ϑ*(*π*) and the number of candidate solutions found. However, in practice, we found that an approach that thresholds the ratio of p-values (default of 5) between the best solution and the next best solution is less conservative and works well for real datasets.

In summary, our statistical scoring approach finds an optimal solution and evaluates its statistical significance and uniqueness in a unified framework, and thus allows for savings in computational time and space compared to a permutation test, without restricting the method to a scenario where experimental error probabilities are known *a priori*.

## 4 Overlap map-to-sequence alignment

In order to extend OPTIMA to compute and evaluate overlap alignments —a key step in assembly pipelines that use mapping data [7,8],— we use a sliding window approach based on OPTIMA. This allows us to continue using the statistical evaluation procedure defined in OPTIMA that relies on learning parameters from comparable alignments —i.e. those based on the same number, size, and order of experimental fragments,— in a setting where the final alignments are not always of the same length and structure. Briefly, for each map, OPTIMA-Overlap first finds optimal sub-map alignments, evaluates their significance and uniqueness, and then tries to extend the candidate alignments found until it reaches the ends of either the experimental map or the *in silico* map, in order to choose the most significant overlap alignments (see Figure 1d). This approach begins by dividing an experimental map into sub-maps of length *l* with a sliding window, and then glocally aligning them to *in silico* maps using OPTIMA (allowing again for truncated ends to account for high error rates). Each glocal alignment sub-problem will then return either:

i. *a significant and unique sub-map alignment;*
ii. *an alignment labeled as non-significant and/or non-unique (which will be considered as false alignments);*
iii. *no feasible alignments found.*

Optimal solutions from the sub-problems are then ranked by p-value (smallest to largest) and iterated through to select sub-maps that should be extended. At each stage we check for significance and uniqueness of the reported solutions (compared to the others) as well potential cases of identical or conflicting alignments, as defined below:

### Definition 3 Conflicting alignments

*A sub-map alignment π*_1_ *is said to be* conflicting *with another alignment π*_2_ *if either*

a. *the sub-map of π*_1_ *overlaps the sub-map of π*_2_, *or*
b. *π*_1_ *aligns to the same* in silico *map of π*_2_*, but in a different location or strand.*

Conflicting alignments may results in ambiguous placement of an experimental map on a database of *in silico* maps, but condition (a) could be relaxed in some cases, e.g. when experimental maps are known to overlap multiple *in silico* maps in the same region. While iterating through the list of sub-maps, the following rules are thus implemented:

1. Significance — if the current solution *π*_*i*_ is labeled as a false alignment, then we stop iterating through the rest of the list.
2. Uniqueness — we skip an alignment if either: (i) *π*_*i*_ represents the same overlap alignment as another more significant solution; (ii) *π*_*i*_ is *conflicting* with a solution having lower p-value (i.e. seen before); or, (iii) *π*_*i*_ is not unique with respect to other solutions *π*_*j*_ with *j* > *i* (i.e. having greater p-values) that it is *conflicting* with.
3. Extension with dynamic programming — optimal solutions according to Equation (2) are identified, where ties are broken in favor of longer alignments.

This approach allows us to report multiple overlap alignments (including split alignments) for an experimental map, while we use the q-value analysis as before to report all alignments with *q ≤* 0.01. In addition, we can reuse the dynamic programming matrix computed for each seed across sub-map alignments and thus complete the overlap alignment with the same asymptotic time and space complexity as the glocal alignment.

## 5 Results and discussion

### Generation of benchmarking datasets

In order to benchmark OPTIMA and OPTIMA-Overlap against other state of-the-art aligners, we first developed synthetic datasets that aim to represent two ends of the spectrum of errors in mapping data for eukaryotic genomes. These scenarios were defined by confidently aligning, using SOMA [14] and manual curation, two sets of maps from different experimental runs for optical mapping on a human cell line. The first scenario, (A), was defined based on lanes that were reported by the Argus machine to have high quality scores, while the second scenario, (B), was defined by lanes with lower quality that were typically obtained on the system. Specifically, we estimated three key parameters from the data: *d*, the restriction enzyme digestion rate, *f*_100_, the false cut rate per 100 kbp, and the fragment sizing errors for predefined *in silico* fragment size ranges (these ware fixed for both scenarios):

A. Easier scenario: *d* = 0.78, *f*_100_ = 0.97, and probability at 50% of missing fragments below 1.2 kbp, at 75% below 600 bp, and at 100% below 350 bp;
B. Harder scenario: *d* = 0.61, *f*_100_ = 1.38, and 50% missing fragments below 2 kbp, 75% below 800 bp, and 100% below 350 bp.

For each scenario, we simulated cut and sizing errors using the probability distributions described in [17] with the above parameters, and map sizes based on empirically-derived distributions from real maps (average size of approximately 275 kbp and containing 17 fragments). We generated 100× coverage of maps with errors sampled uniformly from the *Drosophila melanogaster* (BDGP 5) and *Homo sapiens* (hg19/GRCh37) genomes using the KpnI restriction pattern 

~~~
GGTAC’C
~~~

, which resulted in 13,920 fragments genome-wide (normal and reverse strand) with an average fragment size (AFS) of 17.3 kbp, and 573,276 fragments with AFS=10.8 kbp, respectively.

### OPTIMA results

OPTIMA was compared against the state-of-the-art algorithms Gentig v.2 [2,3,4], SOMA v.2 [14], and Valouev’s Likelihood score [17] for glocally aligning the simulated maps over their respective *in silico* reference genomes. We also ran variations of these algorithms from their default options (d), specifically by providing the true error distribution parameters used in the simulations as input (tp), the adjusted AFS based on the organism under analysis (a), parameter values published in their respective papers instead of the default ones in addition to the true error distribution rates used (p), and by allowing the trimming of map ends in the alignment (t). Moreover, SOMA [14] was modified to correctly handle missing *in silico* fragments up to 2 kbp and to run only for *C*_*σ*_ = 3(v) to make its results comparable. We omitted SOMA’s statistical test (also for Valouev’s Likelihood method) as it is unfeasible for large datasets. TWIN [13] was not included in this comparison as it does not allow for errors and missing information in experimental maps.

As can be seen from the results in Table 1, OPTIMA reports alignments with very high precision, *>*99% in most cases, independent of the genome size and the dataset error rate. In comparison Gentig has similar high precision on the Drosophila genome but lower precision on the human genome, with as low as 80% precision under scenario (B) (with default parameters). Without their computationally expensive statistical tests, which can increase the runtime by a factor *>*100, SOMA and the Likelihood method have much lower precision, particularly on the human genome. In addition, in terms of sensitivity, OPTIMA was found to be notably better than other aligners. In particular, while for the higher quality scenario (A) OPTIMA provides a *>*1.5 × boost over Gentig in sensitivity, for the commonly obtained scenario (B) OPTIMA is more than 2 times as sensitive as Gentig. The relatively high sensitivities of SOMA and the Likelihood-based method in these experiments are likely an artifact of relaxed settings in the absence of their statistical tests. These results highlight OPTIMA’s high precision and improved sensitivity across experimental conditions, and suggest that it could adapt well to other experimental settings as well.

**Table 1:** **Comparison of all methods and their variants on glocal map-to-sequence alignment.** Sensitivity (S) and precision (P) are in percentages and the best values across methods are highlighted in bold. Results are based on the alignment of a subset of 2,100 maps.

| Algorithm | Drosophila (A) |  | Drosophila (B) |  | Human (A) |  | Human (B) |  |
| --- | --- | --- | --- | --- | --- | --- | --- | --- |
|  | S | P | S | P | S | P | S | P |
| OPTIMA | **90** | **100** | **49** | **99** | **83** | **100** | **43** | **98** |
| Gentig v.2 (d) | 59 | **100** | 24 | **99** | 53 | 96 | 20 | 80 |
| Gentig v.2 (tp) | 59 | **100** | 24 | 98 | 54 | 95 | 20 | 88 |
| SOMA v.2 (v) | 72 | 73 | 31 | 39 | 50 | 50 | 17 | 20 |
| Likelihood (d+a) | 49 | 49 | 29 | 30 | 24 | 24 | 14 | 14 |
| Likelihood (d+a+t) | 64 | 65 | 38 | 39 | 33 | 34 | 18 | 19 |
| Likelihood (p+a+t) | 75 | 75 | 39 | 39 | 62 | 62 | 19 | 20 |

In Table 2, we further compare all methods on their running time as well as worst-case complexity (runtime and space). As can be seen here, SOMA and the Likelihood-based methods are at least an order of magnitude slower than OPTIMA and Gentig. Gentig’s proprietary algorithm is based on earlier published work, but its current version uses an unpublished hashing approach. In comparison, OPTIMA is 2 times faster while being *>*50% more sensitive than Gentig, and shows both time and memory complexity improvements over SOMA and Likelihood score. *OPTIMA-Overlap results.* For overlap alignment, we compared OPTIMA-Overlap with an overlap-finding extension of Gentig v.2 (implemented in the commercial software Genome-Builder from OpGen, which contains a module called ScaffoldExtender) [3,4], as well as with Valouev’s Likelihood-Overlap method [17].

**Table 2:** **Running time and worst-case complexity for various glocal map-to-sequence aligners.** Running times reported are estimated from 2,100 maps and extrapolated for the full datasets (82,000 Drosophila maps and 2.1 million Human maps, for 100× coverage; single-core computation on Intel x86 64-bit Linux workstations with 16 GB RAM). The best column-wise running times are reported in bold. Note that including the permutation-based statistical tests for SOMA and the Likelihood method would increase their runtime by a factor >100. The complexity analysis refers to map-to-sequence glocal alignment per map, where *n* is the total length of the *in silico* maps (∼500,000 fragments for the human genome), *m* ≪ *n* is the length of the experimental map in fragments (e.g. 17 fragments), #*seeds*, *c* (default of 2), and *δ* are as defined in Section 3, and #*it* (number of iterations), #*hashes* (geometric hashes found to match), and |*HashTable|* are as partially specified in [3,4].

| Algorithm | Complexity |  | Running time |  |
| --- | --- | --- | --- | --- |
|  | Time | Space | Drosophila | Human |
| OPTIMA | $O((m - c) \#seeds \delta^3)$ | $O((m - c)^2 + cn)$ | **54 m** | **36 days** |
| Gentig v.2 (d) | $O(m \#it \#hashes \delta^3)$ | $O(m^2 + n + \vert HashTable\vert )$ | 1.32 h | 75 days |
| Gentig v.2 (tp) |  |  | 1.85 h | 174 days |
| SOMA v.2 (v) | $O(m^2 n^2)$ | $O(mn)$ | 1.28 years | 1,067 years |
| Likelihood (d+a) | $O(mn \delta^2)$ | $O(mn)$ | 22.22 h | 2.72 years |
| Likelihood (d+a+t) |  |  | 19.62 h | 2.38 years |
| Likelihood (p+a+t) |  |  | 41.73 h | 5.53 years |

In our first test, we randomly selected 1,000 maps for each scenario (A) and (B) from our previously simulated maps for Drosophila and Human genomes. In addition, we simulated assembled sequence fragments (assuming short-read and mate-pair assembly) for Drosophila and Human genomes based on empirical scaffold size distributions (Drosophila assembly N50 of 2.7 Mbp with 239 scaffolds and Human assembly N50 of 3.0 Mbp with 98,987 scaffolds; [18]). Simulated assemblies were then used to generate *in silico* maps (filtered for those with <4 non-end fragments, as these cannot be confidently aligned [1,16]), which were then aligned with the simulated experimental maps.

For our second test, we compared all methods on optical mapping data generated in-house from a human cancer cell line (K562) on OpGen’s Argus System (for runtime reasons, a random sample of 2,000 maps with ≥ 10 fragments was extracted), and *in silico* maps generated from *de novo* assemblies of shotgun Illumina sequencing data (HiSeq) and six mate-pair libraries with insert sizes ranging from 1.1 kbp to 25 kbp [20] (N50=3.1 Mbp, 76,990 scaffolds). This dataset likely represents a harder scenario, with assembly gaps/errors and genomic rearrangements confounding the analysis. It also represents a likely use case where mapping data will be critical to detect large structural variations, disambiguate complex rearrangements, and ultimately assemble cancer genomes *de novo*.

For each test, we evaluated the precision of alignments as well as the number of (correctly) reported alignments that provide an extension to the *in silico* maps through experimentally determined fragments, as this is key for the application of overlap alignments in genome assembly. We begin by noting that there is an important trade-off between sensitivity with a specific window size in OPTIMA-Overlap and the correctness of alignments, as can be seen in Figure 3). As expected, even though small window sizes (less than 10 in Figure 3) provide more sensitive results, they also make true alignments indistinguishable from noise and reduce the number of correct alignments detected. On the other hand, longer window sizes improve the signal-to-noise ratio but lead to a drop in sensitivity, leaving a sweet-spot in the middle (10–13 fragments) where the method is most sensitive across a range of datasets. In particular, real datasets are slightly more challenging than our simulations (Human (B) vs. real data in Figure 3) and so we have conservatively chosen a window size of 12 as the default for OPTIMA-Overlap. By benchmarking OPTIMA-Overlap with this setting we observed similar high precision as observed with OPTIMA for glocal alignment (Table 3). This was seen uniformly across datasets with disparate profiles in terms of genome size and error rates, suggesting that our statistical evaluation is reasonably robust. As before, we also note that Gentig’s approach as well as the Likelihood-based method may not always exhibit high precision. Finally, in terms of sensitivity, OPTIMA-Overlap improves over competing approaches by 30–80%, and this is also seen in the harder real datasets.

**Figure 3:**
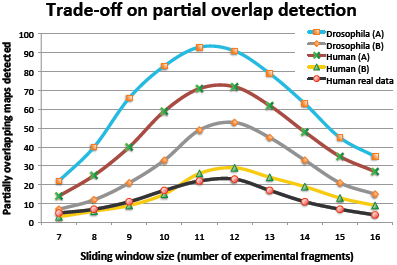
Number of (correct) partial overlaps found for each sliding window size using OPTIMA-Overlap, for both simulated and real maps over simulated and real scaffolds, respectively.

**Table 3:**
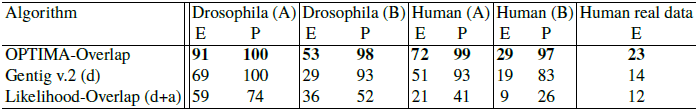
**Comparison of methods for overlap map-to-sequence alignment.** We report the precision of overlap alignments (P, in percentages) and the number of overlap alignments that lead to (correct) extensions (E, absolute values) as a measure of sensitivity (correctness is only known for simulated datasets). The best values across methods are highlighted in bold.

| Algorithm | Drosophila (A) |  | Drosophila (B) |  | Human (A) |  | Human (B) |  | Human real data |
| --- | --- | --- | --- | --- | --- | --- | --- | --- | --- |
|  | E | P | E | P | E | P | E | P |  |
| OPTIMA-Overlap | **91** | **100** | **53** | **98** | **72** | **99** | **29** | **97** | **23** |
| Gentig v.2 (d) | 69 | 100 | 29 | 93 | 51 | 93 | 19 | 83 | 14 |
| Likelihood-Overlap (d+a) | 59 | 74 | 36 | 52 | 21 | 41 | 9 | 26 | 12 |

### Utility in real-world applications

Overlap alignments form a critical building block for applications such as OpGen’s Genome-Builder and its use in boosting assembly quality [7]. As OPTIMA-Overlap can work with lower quality data (scenario (B) in our simulations; Genome-Builder would typically filter out such data) and also provides improved sensitivity in detecting overlap alignments, we estimated that its use could reduce the requirement for generating mapping data by half. As the cost of mapping data for the assembly of large eukaryotic genomes can range from USD 20,000 to 100,000, this can lead to significant cost savings.

Similarly, we tested OPTIMA and Gentig on data for a human cell line (HCT116) generated on two runs of the Argus System from OpGen [7], in order to calculate how much mapping data would be needed for sufficient aligned coverage of the human genome to enable structural variation analysis. Using two sets of 31,588 and 36,018 maps, respectively, that were aligned to the human reference genome, we found that OPTIMA confidently aligned 42% of the maps vs. 26% by Gentig (with default parameters) for the first (easier) dataset (1.6× increase), and 17% of maps vs. 9% for Gentig in the second (harder, but more commonly obtained) dataset (1.9× increase). These results suggest that for structural variation analysis on the human genome, particularly for cancer genomes, OPTIMA can reduce mapping costs by 38–47%, thus saving tens of thousands of dollars in project cost as well as enabling faster analyses of the data.

## 6 Conclusion

With the availability of new mapping technologies (e.g. Nabsys) and greater use of existing ones to complement highthroughput sequencing, there is a critical need for robust, publicly-available computational tools that can combine mapping and sequence data efficiently. In this work, we introduce two new alignment tools that address this need for a wide range of applications from genome assembly to structural variation analysis. Our benchmarking results provide evidence that these methods outperform existing approaches in sensitivity and runtime while providing highly precise alignments in a range of experimental settings. Similar results were also seen in real datasets from human cell lines, suggesting that they could help in significantly reducing the cost of optical mapping analysis needed, and thus increase its usage as well.

In the development of OPTIMA and OPTIMA-Overlap we establish two key new ideas for map-to-sequence alignment. The first is the introduction of composite seeds, an idea that echoes the idea of spaced seeds in the context of continuous-valued sequence alignment. Composite seeds allowed us to develop efficient seed-and-extend aligners for map-to-sequence alignment of highly erroneous mapping data. We believe that similar ideas can also be applied for map-to-map alignment and *de novo* assembly of experimental maps. The second concept is the development of a statistical testing approach that does not require knowledge about the true distribution of errors, or an expensive permutation test to evaluate the uniqueness and significance of alignments. This allowed us to significantly reduce the runtime cost of this step, without sacrificing specificity or the ability to be agnostic to error rates. While our experiments with real data in this work were limited to data generated on the Argus System from OpGen, similar ideas (with minor variations) should also be applicable to data from other technologies such as the Irys Platform from BioNano Genomics.

In future work, we plan to implement further runtime and memory optimizations to OPTIMA and OPTIMA-Overlap and explore their use for super-scaffolding of large genomes [18], as well as for studying genomic rearrangements in cancer.

## Availability

http://www.davideverzotto.it/research/OPTIMA

## References

1 Anantharaman, T.S., Mishra, B.: A probabilistic analysis of false positives in optical map alignment and validation. In: First International Workshop on Algorithms in Bioinformatics. pp. 27–40. WABI 2001, Springer (2001)

2 Anantharaman, T.S., Mishra, B., Schwartz, D.C.: Genomics via Optical Mapping II: Ordered restriction maps. Journal of Computational Biology 4(2), 91–118 (1997)

3 Anantharaman, T.S., Mishra, B., Schwartz, D.C.: Genomics via Optical Mapping III: Contiging genomic DNA. In: ISMB 1999. pp. 18–27. AAAI (1999)

4 Anantharaman, T.S., Mysore, V., Mishra, B.: Fast and cheap genome wide haplotype construction via optical mapping. In: Altman, R.B., Jung, T.A., Klein, T.E., Dunker, A.K., Hunter, L. (eds.) Pacific Symposium on Biocomputing 2005. World Scientific (2005)

5 Antoniotti, M., Anantharaman, T., Paxia, S., Mishra, B.: Genomics via Optical Mapping IV: Sequence validation via optical map matching. Tech. rep., New York, NY, USA (2001)

6 Califano, A., Rigoutsos, I.: FLASH: a fast look-up algorithm for string homology. In: ISMB 1993. pp. 56–64. AAAI (1993)

7 Dong, Y., Xie, M., Jiang, Y., Xiao, N., Wang, W., et al. : Sequencing and automated whole-genome optical mapping of the genome of a domestic goat (*capra* hircus). Nature Biotechnology 31, 135–141 (2013)

8 Ganapathy, G., Howard, J., Ward, J., Li, J., Phillippy, A., Jarvis, E., et al: High-coverage sequencing and annotated assemblies of the budgerigar genome. GigaScience 3(1), 11 (2014)

9 Karp, R.M., Rabin, M.O.: Efficient randomized pattern-matching algorithms. IBM J. Res. Dev. 31(2), 249–260 (1987)

10 Kenny, D.A.: Statistics for the Social and Behavioral Sciences. Little, Brown (1987)

11 Lam, E.T., Hastie, A., Lin, C., Ehrlich, D., Nagarajan, N., Xiao, M., Kwok, P.Y., et al. : Genome mapping on nanochannel arrays for structural variation analysis and sequence assembly. Nature Biotechnology 30(8), 771–776 (2012)

12 Lin, H., Goldstein, S., Mendelowitz, L., Zhou, S., Wetzel, J., Schwartz, D., Pop, M.: AGORA: assembly guided by optical restriction alignment. BMC Bioinformatics 13(1), 189 (2012)

13 Muggli, M.D., Puglisi, S.J., Boucher, C.: Efficient indexed alignment of contigs to optical maps. In: Algorithms in Bioinformatics, LNCS, vol. 8701, pp. 68–81. Springer (2014)

14 Nagarajan, N., Read, T.D., Pop, M.: Scaffolding and validation of bacterial genome assemblies using optical restriction maps. Bioinformatics 24(10), 1229–1235 (2008)

15 Ray, M., Goldstein, S., Zhou, S., Potamousis, K., Schwartz, D., et al. : Discovery of structural alterations in solid tumor oligodendroglioma by single molecule analysis. BMC Genomics 14(1), 505 (2013)

16 Sarkar, D., Goldstein, S., Schwartz, D.C., Newton, M.A.: Statistical significance of optical map alignments. Journal of Computational Biology 19(5), 478–492 (2012)

17 Valouev, A., Li, L., Liu, Y.C., Schwartz, D.C., Yang, Y., Zhang, Y., Waterman, M.S.: Alignment of optical maps. Journal of Computational Biology 13(2), 442–462 (2006)

18 Verzotto, D., Hillmer, A.M., Teo, A.S., Nagarajan, N.: Super-scaffolding of large eukaryotic genomes with single molecule landmark maps (2015), manuscript in preparation

19 Wilson, E.B., Hilferty, M.M.: The distribution of Chi-Square. PNAS 17(12), 684–688 (1931)

20 Yao, F., Ariyaratne, P.N., Hillmer, A.M., Liu, E.T., Ruan, Y.: Long span DNA paired-end-tag (DNA-PET) sequencing strategy for the interrogation of genomic structural mutations and fusion-point-guided reconstruction of amplicons. PLoS ONE 7(09), e46152 (2012)

